# Neuropilin-1 Assists SARS-CoV-2 Infection by Stimulating the Separation of Spike Protein Domains S1 and S2

**DOI:** 10.1101/2021.01.06.425627

**Authors:** Zhen-lu Li, Matthias Buck

## Abstract

The cell surface receptor Neuropilin-1 (Nrp1) was recently identified as a host factor for SARS-CoV-2 entry. As the Spike protein of SARS-CoV-2 is cleaved into the S1 and the S2 domain by furin protease, Nrp1 binds to the newly created C-terminal RRAR amino acid sequence of the S1 domain. In this study, we model the association of a Nrp1 (a2-b1-b2) protein with the Spike protein computationally and analyze the topological constraints in the SARS-CoV-2 Spike protein for binding with Nrp1 and ACE2. Importantly, we study the exit mechanism of S2 from the S1 domain with the assistance of ACE2 as well as Nrp1 by molecular dynamics pulling simulations. In the presence of Nrp1, by binding the S1 more strongly to the host membrane, there is a high probability of S2 being pulled out, rather than S1 being stretched. Thus, Nrp1 binding could stimulate the exit of S2 from the S1 domain, which will likely increase virus infectivity as the liberated S2 domain mediates the fusion of virus and host membranes. Understanding of such a Nrp1-assisted viral infection opens the gate for the generation of protein-protein inhibitors, such as antibodies, which could attenuate the infection mechanism and protect certain cells in a future combination therapy.

## Main Text

The virus SARS-CoV-2 responsible for the world-wide 2020/21 COVID-19 pandemic, is also increasingly appreciated for its deleterious effects on the human nervous and cardiovascular systems, including longer term symptoms in some of the patients. While intersecting with both system, the VEGFR2 and plexin co-receptor, Neuropilin-1 was also recently identified as a host factor for SARS-CoV-2.^1-3^ Following binding to the primary extracellular receptor, the ACE2 protein at the Receptor Binding Domain (RBD), -but possibly independent of it-the Spike protein is cleaved via a host cell protease Furin, which appears to be a crucial step of SARS-CoV-2 for cell entry.^4-5^ This type of cleavage generates two protein fragments, S1 and S2, of which the former exposes a newly created C-terminal polypeptide chain with the sequence RRAR.^6,7^ This amino acid motif allows the S1 protein to interact with other cell surface receptors, such as Neuropilins (Nrps).^1-3,8^ Eventually S1 and S2 separate, with S2 displaying a membrane fusion domain (which was previously largely hidden by S1 in the structure of the virus Spike protein). Then S2 domain triggered fusion allows the flow of virus RNA into the host cell for replication, protein synthesis and new virus assembly.^4^ The newly generated S1 C-terminal poly-arginine sequence is recognized by the b1 domain of Nrp1, the same site used by VEGF ligands to interact with Nrp1.^8-10^ This suggests that Spike protein binding will likely block VEGF-Nrp1 interactions as they occur in normal cells. It will also interfere with endogenous receptor complex formation and endogenous signaling/functions in the host cell involving Nrp1 either directly or indirectly.^11-13^ Due to involvement of Neuropilin receptor in the development and function of axons and synapses, and VEGF in the growth/leakiness of blood vessels, the SARS-CoV-2 infection may have long lasting effects on the nervous and cardiovascular systems.^14-16^ Specifically, a significant number of critically ill COVID-19 patients experience impairments such as brain inflammation, an increased permeability of the blood brain barrier as well as an elevated occurrence in the number of strokes.^17-20^ These observations make the Nrp1 – SARS-CoV-2 interaction an urgent focus for investigation.

It is now well known that the initial contact of SARS-CoV-2 is made with the human ACE2 receptor on the host cell via the prominent extracellular feature of coronaviruses, the Spike protein.^21^ However, it is still unclear whether Nrp1 may be a co-binding factor together with ACE2 and how, if at all, the two interact; either directly or indirectly. Moreover, while the S1 C-terminal RRAR segment is a major recognition site for Nrp1, it is not yet known which other regions of Nrp1 and the Spike protein may interact. Experimentally, Nrp1 was found to facilitate the virus infection and even interacts with the Spike protein when the RRAR segment is deleted.^1-3^ However, the details of the infection mechanism -and the role of Nrp1 in it-are not yet understood. In this study, we predict possible binding models between a Nrp1 a2-b1-b2 domain containing protein with the ACE2 bound Spike protein (Fig. 1). By considering topological constraints, our docking calculations identify probable binding models of Nrp1 with the Spike protein trimer. Importantly, molecular dynamics pulling simulations suggest that the exit mechanism of S2 from the S1 domain is assisted by membrane bound Nrp1, defining a role for Nrp1 in the SARS-COV-2 in modulating separation of the Spike protein S1 and S2.

**Figure 1:**
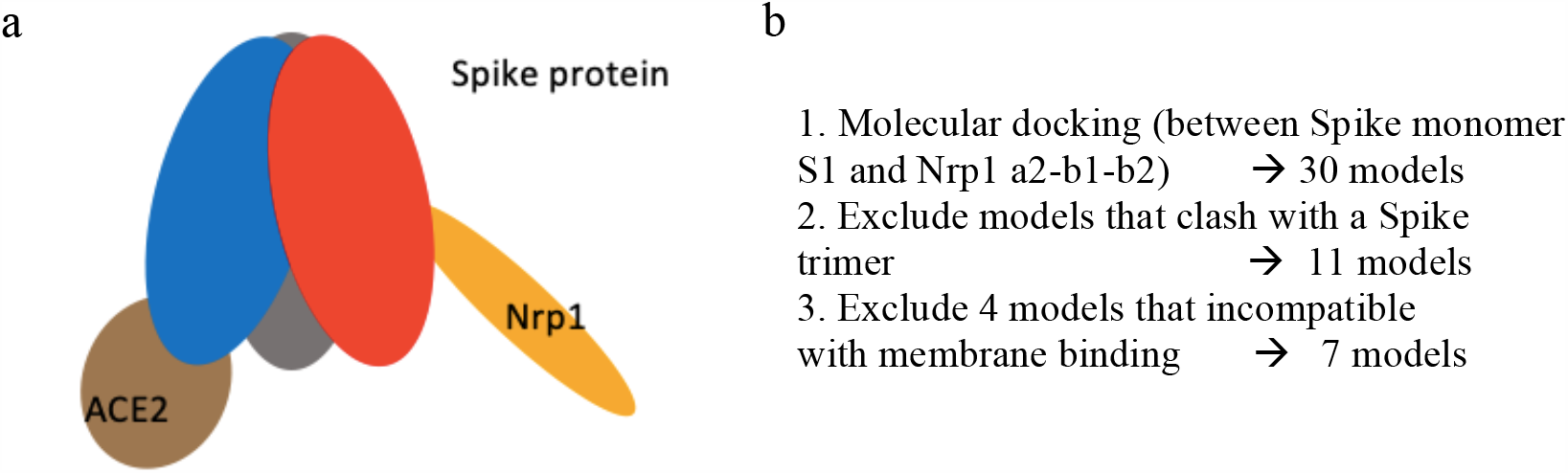
(a) Schematic Model of Spike Protein trimer bound to one ACE2 and one Nrp1 receptor. (b) Molecular docking yields 7 possible binding models which satisfy geometric constraints.

## Results and Discussion

### Modeling of Nrp1: Spike protein complexes

The polybasic sequence 678-TNSPRRAR-685, created by cleavage of furin of the Spike protein as the new C-terminus of the Spike protein (called S1 domain), was shown to associate with the b1 domain of Nrp1.^1^ We firstly modeled a complex of a Nrp1 a2-b1-b2 domain containing protein bound to this polybasic peptide based on the template of the Nrp1 a2-b1-b2: VEGF-A complex (PDB entry 4deq).^10^ The modeled complex was set up and then relaxed for 20 ns at 310 K by an all-atom simulation as described in the Methods section. The association between the S1 C-terminal peptide and Nrp1 b1 domain is largely maintained by electrostatic attraction of Spike S1 residues R683 and R685 with Nrp1 residues D320 and D348 as characterized in the study of Daly et al.^1^ While the RRAR segment is a major recognition site for Nrp1, the study also presented evidence that other Nrp1 regions may interact with the Spike protein.^1^ In order to identify such sites, we docked the relaxed structure of the simulated Nrp1-peptide complex onto a Spike protein monomer (residues 1 to 677) with ClusPro 2.0.^22^ The structure of the Spike protein, which was based on available crystal structures and models (adding missing loops and residues) has been developed and made available by Woo et at.^23,24^ For simplicity, glycosylation of the Spike protein was not considered in the modeling. In the docking, distance restraints were applied between two residues: the last residue of Spike protein monomer Q677 and the first amino acid of the polybasic peptide T678 (further details are given in the Methods section). The molecular docking via ClusPro gave 30 predicted structures. First, we merged similar models and excluded models where the Nrp1 a2-b1-b2 domain would clash with any region of a full-length Spike trimer (residues 1-1146). This results in 11 models which are still compliant and are further discussed below. We excluded 4 other models due to constraints set up by the proteins in their distance and orientation to the membrane. We will explain this point below with respect to Fig. 5. So here we considered 7 models for further analysis (Fig. 1b).

### Binding modes of between Nrp1 and Spike protein

We analyze the 7 models which are compatible with the topological constraints (avoiding clashes with regions in one trimer unit (Fig. 2a) and with the other two timer units and the membrane, see Fig. 5 & below). In order to relax and better accommodate the initial docked structures in the context of the full length (1-1146) Spike protein trimer, we performed all atom molecular dynamics simulation of the Spike protein trimers with one bound Nrp1 (a2-b1-b2) for each of the 7 models for 20 ns at 310 K. The experimental study of Daly et al suggested that in addition to the RRAR motif, a second binding site may exist between NTD and RBD regions.^1^ In all 7 models, there are additional interaction beyond binding the S1 C-terminal RRAR motif. These interactions are established between the Nrp1 a2-b1-b2 domain protein with the Spike protein N-terminal (13-305, NTD) and/or the N- and C-termini of the Receptor Binding Domain (RBD, 318-541) or the C-terminal S1 regions (see Table S1 for details). Intriguingly, in our models Nrp1 domains do not interact with the RBD’s motif for binding ACE2 (a sub-region comprising residues 437-538), thus the ACE2 association with the Spike protein should be unperturbed unless there are allosteric effects.^25-27^ The Nrp1 a2-b1-b2 domains also do not interact extensively with S2 (just the very N-terminus in case of models 3,5 and 6) as it would make the Nrp1 domains too distant from the membrane (see below & Fig. 5). In Fig. 2b, we list the principal binding sites (as residue contact x: y in the Spike: Nrp1 complex) for the different models. In model 1, 2 and 7, noticeable hydrophobic contacts such as V622: W411, I68/V70: M204 or I624/R634:Y322/W411 are seen. In model 3, 4, 7, electrostatic pairs such as R21: E456, R214: E308 or R78: E507 are apparent. For the remaining contacts, they are mostly established between polar amino acids with other polar/charged/aromatic amino acid sidechains, which have the possibility to form hydrogen bonds. In supplementary Fig. S1 residues are displayed, which are involved in S1:NRP1 interfaces at least 30% of the time and with a wider cut-off of 0.6 nm. Several models overlap with respect to at least some of the regions which the proteins involved in forming a complex. Intriguingly there is only one model which has all three Nrp1 domains involved and one with only b1 involved (models 4 and 5 respectively). Models involving Nrp1 domains a2, b1 or b1, b2 are considerably varied in the regions which are being contacted on the side of the spike protein. However, for all the seven models, the Nrp1 domains are located in between the NTD (res. 13-305) and the bottom of the RBD (res. 318-330 and 525-541) of the Spike protein. The Nrp1 domains as well as the Spike protein binding regions are relatively extended, with only 3 pairs of regions of the 12 residues defined in Fig. S1, being in close proximity (N2-N4, RBD^N^-RBC^C^ and C2-C3). Interestingly, on the side of the RBD, the RBD of another Spike protein unit would come in depending on whether this RBD was “up” or “down” and it is therefore possible that Nrp1 binding has an effect on this equilibrium. This issue, as well as the interactions that the RBDs make with S2 – possibly facilitating the separation of S1 from S2 requires further investigation beyond the scope of this initial report.^28,29^ Irrespective of such issues, we note that similar to other dynamic protein complexes (e.g. K-Ras at membrane),^30,31^ the nature of the Spike protein: Nrp1 complex suggests that several binding interfaces can interconvert in their use and their occupancies can be transient. The latter is also reflected, in comparison to reports of the SARS-CoV-2 RBD: ACE2 interaction energy^32^ in wider variation of atom-atom pairwise interaction energies between Nrp1 a2-b1-b2 and the Spike protein, as well as the complex-buried area of the accessible surface between the models. A full analysis of the energy landscape and estimation of free energies of binding is a computationally exhaustive endeavor and beyond the scope of this report. Nevertheless, our analysis suggests that rather than forming a single or very few well defined complex structures with specific domain-domain interactions, the purpose of the interactions (which are additional to those of the cleavage generated S1 C-terminal peptide) is to strengthen this principal interaction between the Nrp1 and S1 protein. As discussed below, the other function of the S1 interactions with the somewhat flexible multi-domain Nrp1 is to provide an additional anchor point to the lipid bilayer.

**Figure 2:**
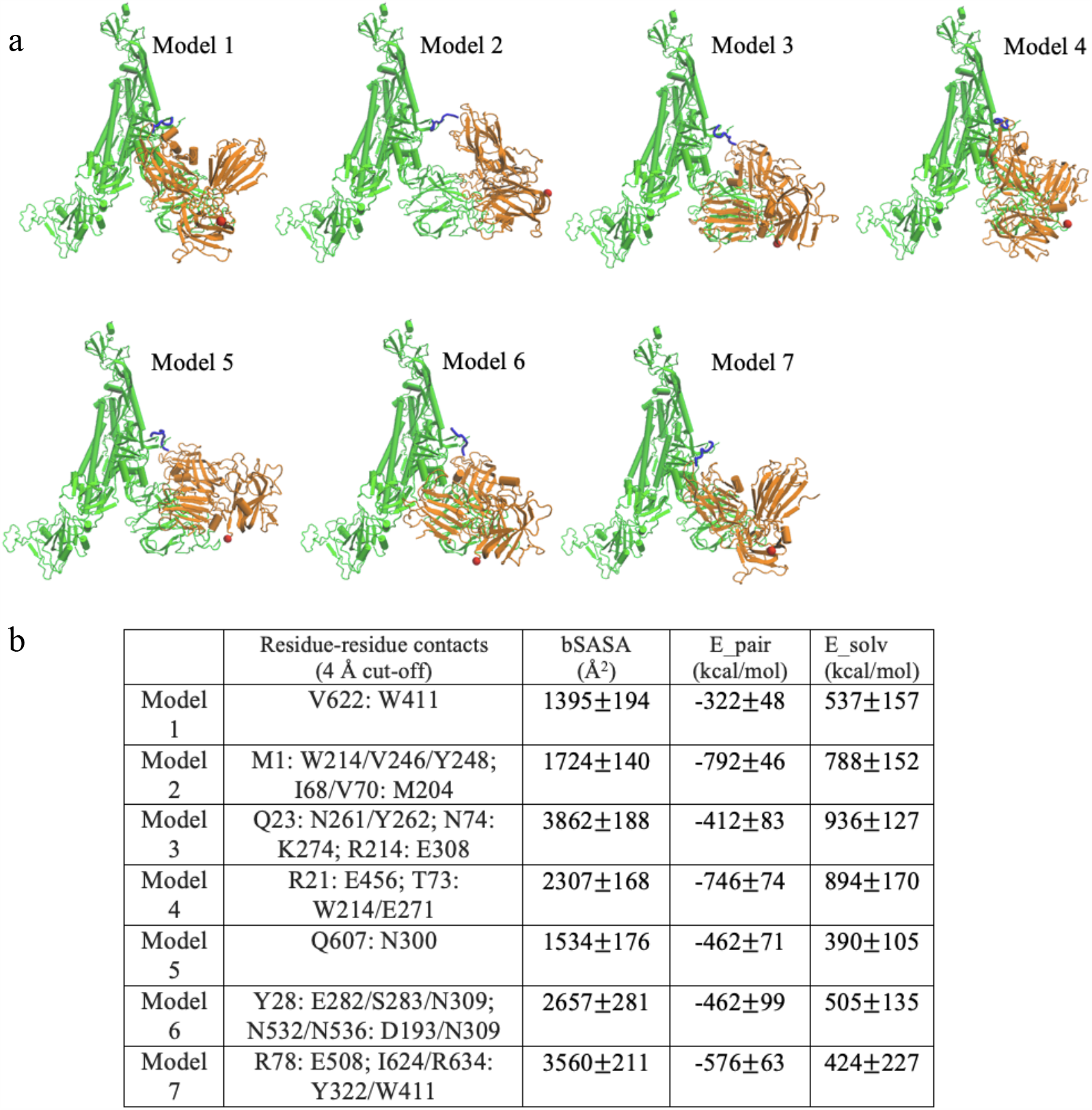
Binding modes for 7 predicted models. (a) Models, shown for Chain C: Nrp1 a2-b1-b2 binding. The TNSPRRAR motif is marked in blue. The last residue of Nrp1 a2-b1-b2 domain is marked in red. (b) Noticeable residue-residue contacts (< 0.4 nm) between Spike protein and Nrp1 in different models (except the RRAR binding region). Together with residue-residue contacts (< 0.6 nm) the proteins are seen to interact via domains a2,b1 and b2 on Nrp1 and a limited number of regions N1-N4, RBD^N^, RBD^C^, C1-C3, the very C-term. of S1 and beginning of S2 (see Table S1 for details). Buried SASA, pair interaction energy and changes of solvation energy for different models. The interaction energy (based upon pairwise vdW and electrostatic forces) are calculated between Nrp1 a2-b1-b2 with a full Spike trimer (Chain A, B, C). The models shown are listed in order of their population near cluster centers (models) 1 to 7 (most to least).

At this point, experimentally there is no binding affinity characterization as well as structural characterization of Nrp1 a2-b1-b2 association available with a SARS-CoV-2 Spike protein. Here our docking/modeling structures could provide reference information for the topology and energetics of the interaction and suggest likely hotspot residues for possible associations between Nrp1 a2-b1-b2 with the Spike protein. More importantly, with the knowledge of possible association models between Nrp1 a2-b1-b2 with the Spike protein, we could examine the exit mechanism of S2 domain out of S1 pocket with the assistance of Nrp1.

### Exit of S2 from the S1 domain

We explored the separation process of the Spike protein by pulling the S2 domain out of the S1 domain in a molecular dynamics pulling simulation of 20 ns. The peptide bond between residues 685 (S1) and 686 (S2) were cut for all three units of the Spike protein trimer. We compared two kinds of complexes: the Spike protein: ACE2 complex and Nrp1: Spike protein: ACE2 complex. Fig. 3 shows representative snapshots of S2 exit from S1 domain with/without the Nrp1 binding. Remarkably, the separation of S2 and S1 for Chain C of the Spike protein happens at earlier steps with Nrp1 bound (∼7 nm versus 16 nm in Fig. 3). The separation of S2 from S1 involves two steps. For the first step, the S2 exits from a cap which consists of S1 domain residues (Fig. 4a). This involves the breaking of the majority of contact interactions between S1 and S2. Residues of S1 RBD (318-541) which interact with the S2 (686-1146) are R319, P322, G381, V382, S383, T385, K386, L390, P412, G413, D427, T430, and L517. Actually, ACE binding appears to require at least one of RBD to be in “up” conformation, which reduce the interaction between one RBD unit and S2.^33^ After step 1 separation, there almost no contacts between S2 and S1 RBD (318-541) remain, however, there is still connection at the cleavage site (685/686) (Fig. 4a, b). Noticeably, S2 residues 692-697 forms a beta sheet with S1 residues 669 to 674. Because of this interaction (Fig. 4b), without the Nrp1 binding, the S1 C-terminal region would tend to drift in together with the S2 domain during separation. This likely accounts for the second barrier during the S1: S2 dissociation and is unfavorable for the separation of the S2 and S1 domains in step 2 (Fig. 4b). Overall, step 1 involves a displacement of S2 of 5 nm, but it has large energy barrier (Fig. 4c). Step 2 involves a longer process of displacement of S2 with a lesser energy barrier. The binding of 682-RRAR-685 motif with Nrp1 at the cleavage site provides a mechanical support for the adjacent regions (∼ S1 res. 610-685). This appears to largely restrict the stretching/local unfolding of the S1 C-terminal region, destabilizes and accelerates the separation of N-terminal segment of S2 domain (∼ res. 686-700) from this region (Fig. 4a).

**Figure 3:**
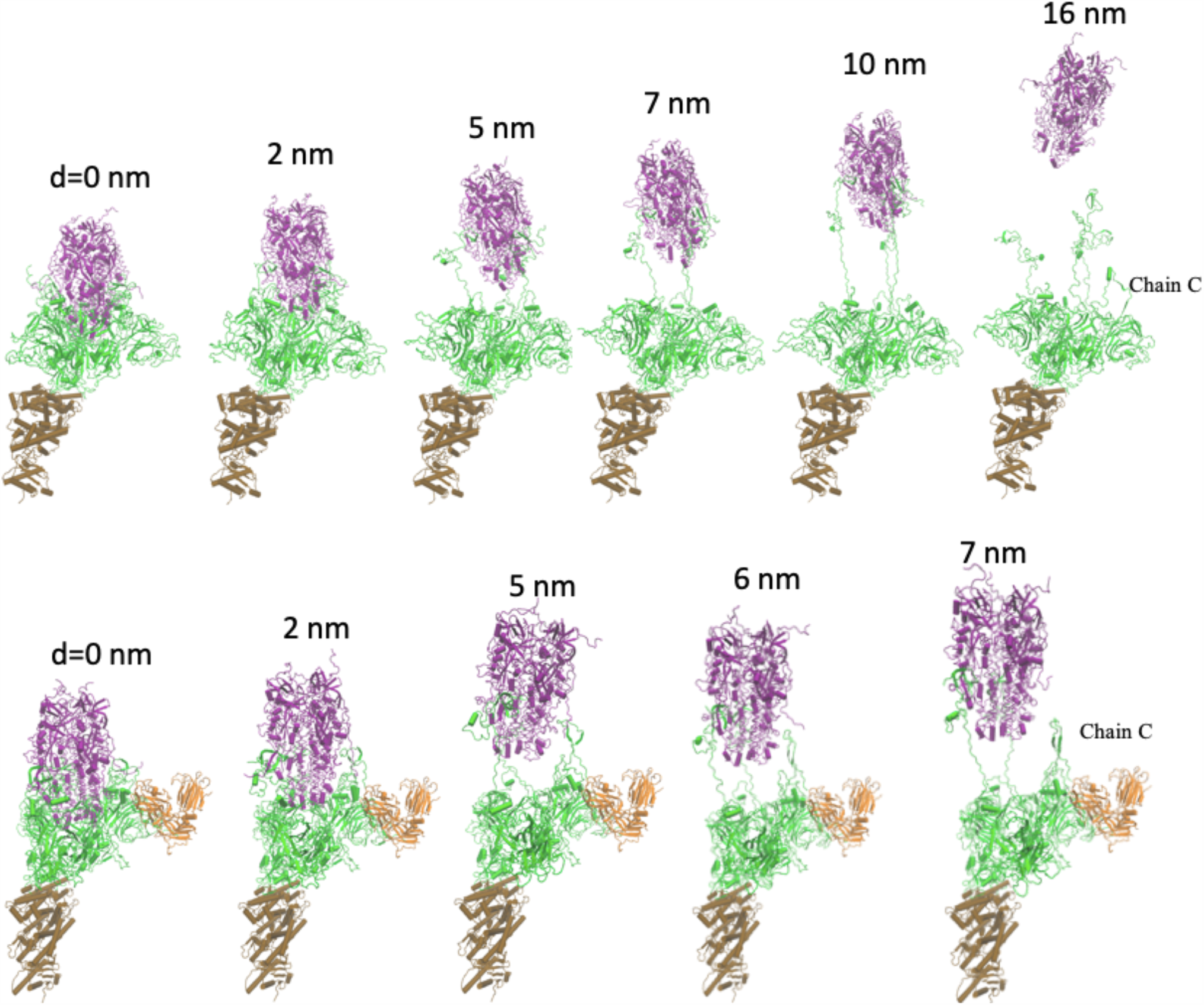
Separation process of S2 and S1 domain of Chain C with/without Nrp1; here shown in absence of Nrp1, top (1st. of 7 simulations shown) and for Nrp1: Spike model 7, bottom. (Color coding: S1 in green and S2 in purple; Nrp1 in orange) The displacement of S2 domain center of mass relative to initial position is given as distance d. The time interval between the structures is 2, 5, 7, 10, 16 ns or 2, 5, 6, 7 ns in the pulling simulation (1 nm/ns).

**Figure 4:**
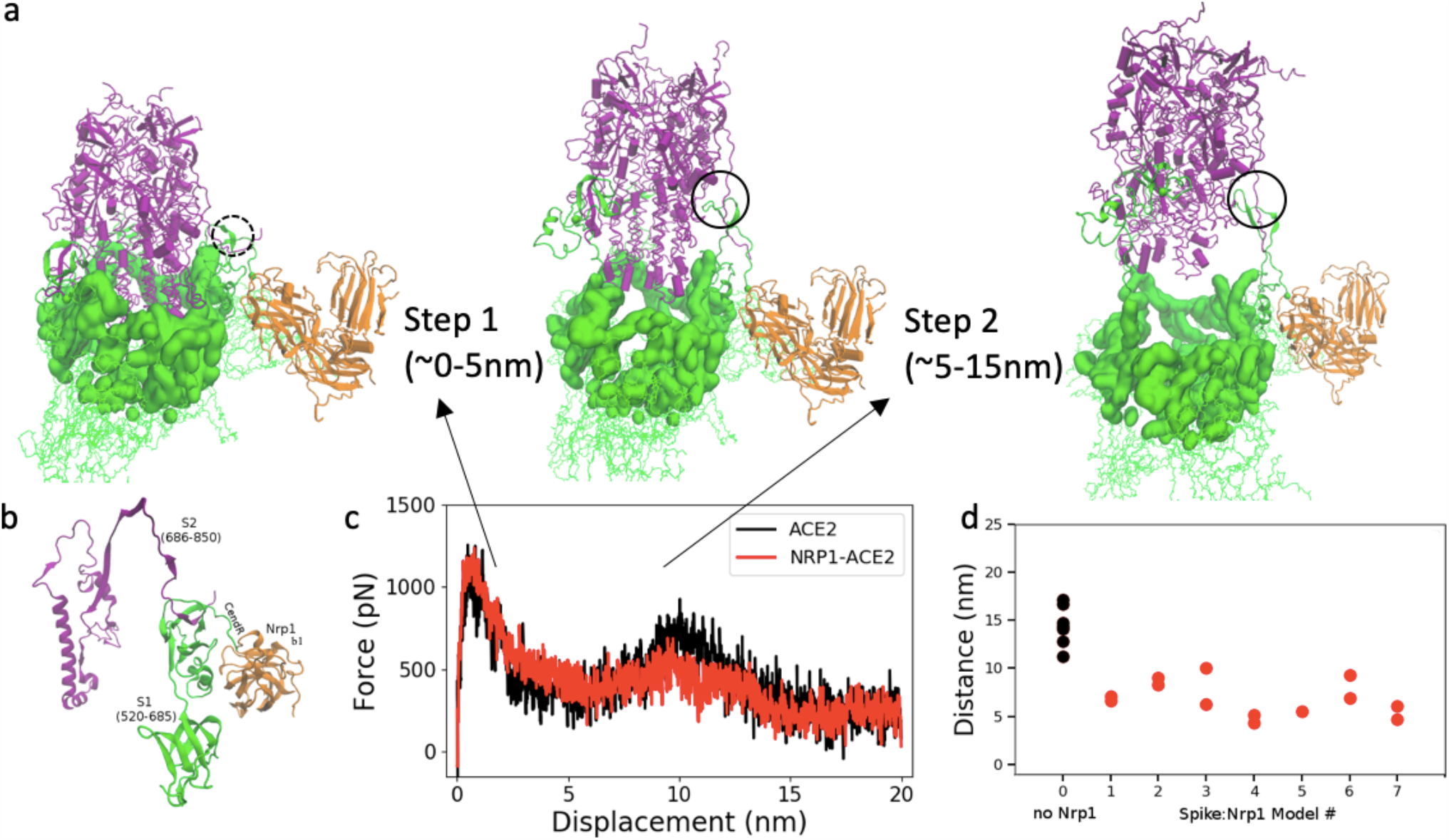
(a) Two steps of S2 exit from S1 domain: S1 regions that cap S2 domain (purple) are highlighted in surface representation in green. (b) S1:S2 contacts at the cleavage site at the start of simulation (model 7 at 0 ns). (c) Pulling force versus displacement of S2 domain (with and without Nrp1). Averaged over simulations of 7 models for simulations without and with Nrp1. (d) Distance between S1 and S2 centers of mass of Chain C of the Spike protein at the time of separation (nearest distance >5Å) between the two domains (of chain C). Two repeats were done for each model except model 5, where Nrp1:S1 binding is easily disconnected during the pulling of S2 domain.

Fig. 4d shows the separation position of S2 out of S1 for chain C of the Spike protein. The separation position is measured by the displacement of center of mass (c.o.m) of S2 to the initial position. For all seven simulations, the separation of S2 and S1 for Chain C of Spike protein happens with a shorter displacement of S2 domain. In models 1-7, it is obvious as the displacement of S2 domain is typically less than 10 nm, mostly around 5-7 nm with the assistance of Nrp1 binding. In contrast, without the Nrp1 binding, the displacement of S2 domain is around 14.4±2.0 nm. So far we have focused on Nrp1 binding with Chain C of Spike protein. Apparently, the Nrp1 binding stimulates the separation of S2 and S1 for Chain C of the Spike protein. However, for Chain A and Chain B, the separation of S2 and S1 is not influenced by Nrp1 binding with Chain C. Typically, the separation of S2 from S1 for Chain A and Chain B needs a displacement of S2 domain about 13-17 nm. Comparing the work needed to separate S2 and S1, the value with/-out Nrp1 is 1113±85 kcal/mol and 1178±91 kcal/mol respectively. There is very little difference between them as the Nrp1 only stimulates the separation of S2 and S1 for Chain C but not for the entire Spike protein trimer.

Having established differences with the ease of the Spike S1:S2 domain separation process, upon Nrp1 binding, we further analyzed the overall structures of the complex at the membrane (Fig. 5a).

**Figure 5:**
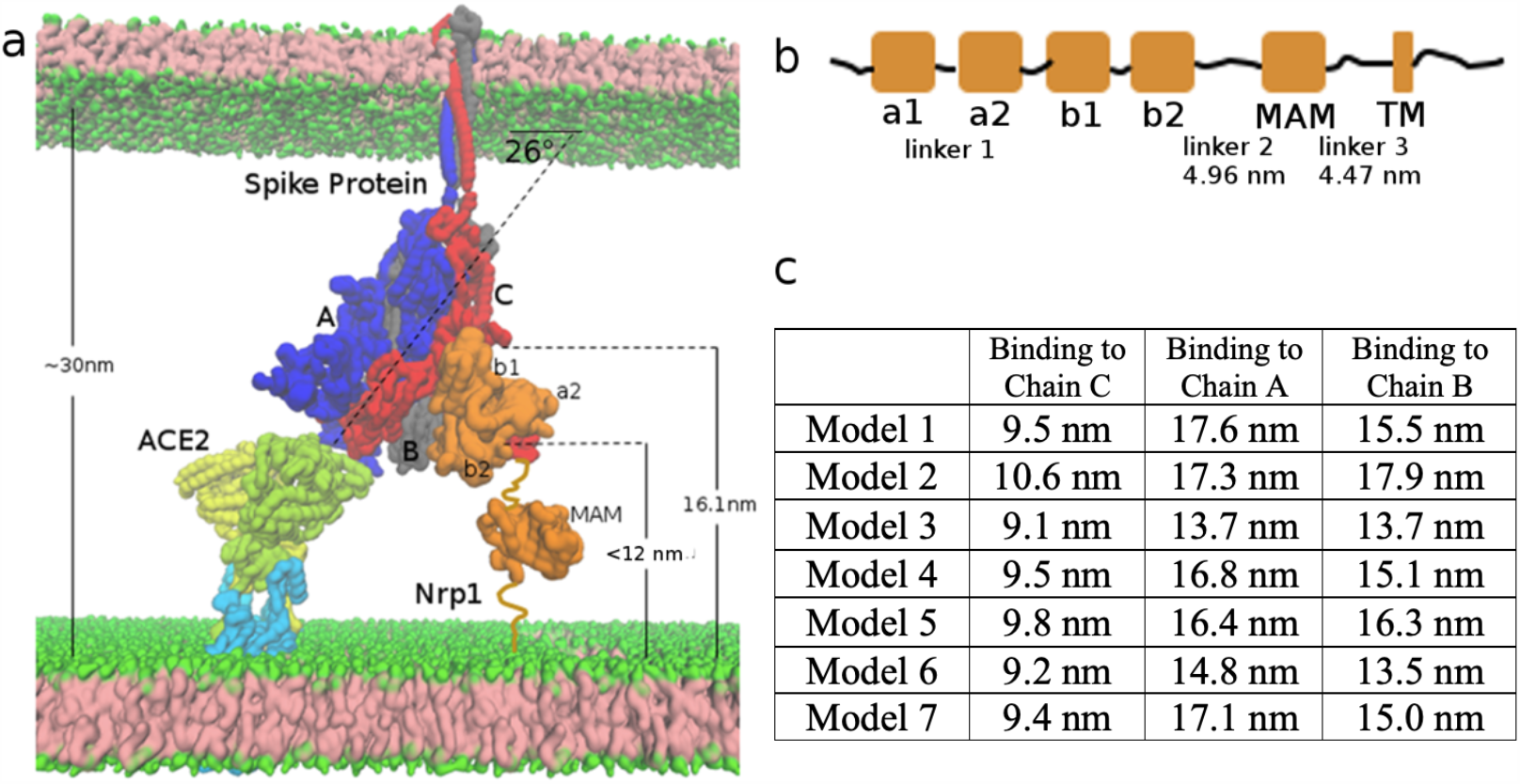
Topological constraints of Nrp1 on binding with Spike protein trimer, bound to an ACE2 dimer. (a) ACE2 (yellow and limone, which also go into and through the membrane) and B^0^AT1(light blue) 2:2 dimer complex at the bottom left. Spike protein trimer (Chain A, B, C in blue, grey and red) in the middle. Nrp1 (orange) at the bottom right of the figure. (b) Linear domain arrangement in the sequence of Nrp1 receptor. (c) Estimated distance of C-terminus of the b2 domain to the membrane bilayer lipid headgroups in the different models with Nrp1 bound to Chain A, B, C.

### Global topological constraints in a Nrp1/ Spike/ACE2 protein complex

Here we consider models of Nrp1 binding to the Spike protein when it is bound to ACE2. Similar to ACE2, Nrp1 is a transmembrane receptor.^8^ The extracellular region (res. 1-856) consists of five sub-domains: a1 (27-141); a2 (147-265); b1 (275-424); b2 (431-586) and a MAM domain (640-813) (Fig. 5b). The fold of the protein fragment containing the a2-b1-b2 domains is relatively compact with short linkers of 5-9 amino acids between domains. By contrast, to the connection between the a1 and a2 domains (linker 1, res. 142 to 146), linkers between b2 and the MAM domain (linker 2, 587-639) as well as the connection of MAM domain to the transmembrane helix (linker 3, 814-856) are long and flexible. The MAM domain crystal structure has been reported.^34^ The distance between the N- and C-terminal residues of the MAM domain (res. 640-813) is about 2.6 nm. As Nrp1 acts as an enhancer for virus infection, we hypothesize that Nrp1 could work as a cofactor that binds to Spike protein together with ACE2. In Fig. 5a, we build a system of an ACE2 bound to Chain A of the Spike protein trimer. ACE2 forms dimer at the membrane surface in complex with the neutral amino acid transporter B^0^AT1 dimer,^35^ Overall, the configuration of the ACE2 dimer lacks flexibility and is perpendicular to the cell membrane, i.e. facing away from it. When bound to ACE2 via a RBD binding interface, the S1-S2 domains are tilted relative to the membrane with a tilt angle of about 26° (orientation of the vector connecting centers of mass (c.o.m) of S1 (1-685) and c.o.m. of S2 (686-1146)). Actually, even without ACE2 binding, the recent cryo-ET structure of an isolated SARS-CoV-2 virus shows a prevalent tilting of the entire Spike protein by 40° relative to the virus surface.^36^ Thus, overall these tilts are compatible with the association between the spike RBD and ACE2 interfaces. While it is also geometrically possible for a spike protein trimer to be bound simultaneously by two different ACE2 receptor dimers, a recent study found that such an interaction involves little avidity effect (i.e. little additional affinity appears to be generated).^37^

As the Spike protein is tilted when it interacts with one ACE2 binding region, Chain C of the Spike protein trimer is the closest subunit to the host membrane. Thus, we consider the binding of Nrp1 to one Spike protein, here Chain C, first. In this construct, ACE2 accesses to the “bottom” RBD domain (Chain A) of the Spike protein trimer; (for binding additional ACE2 and Nrp1, see below). It should be noted that the Nrp1 binding site (the TNSPRRAR motif), in part due to the tilt and position of Chain C has some distance from the host membrane. In fact, the distance of residue Q677 of chain C to the cell membrane surface, as measured in our model of Fig. 5a, is around 16.1 nm. We estimate the end-to-end distance of linker 2 and linker 3 by applying the free rotation model of a polymer chain: 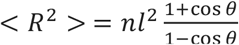 with n as 53 and 43 segments (n) respectively.^38^ The length (*l*) of a segment is set as 0.34 nm and the rotation angle (θ) is set as 134° as typically used in the coarse-grained model for a random coil polypeptide chain.^39^ The average end-to-end distance is estimated as 4.96 nm and 4.47 nm respectively (Fig. 5b). Based on the available structure, the distance between N- and C-terminus of MAM domain is 2.6 nm. Supposing linker 2, linker 3 and MAM all align in a way to maximize the distance in membrane normal direction (Z direction), we estimate the probable length of linker 2-MAM-linker 3 region is 12.0 nm. Therefore, in binding of the Nrp1 a2-b1-b2 with the TNSPRRAR motif, the distance of C-terminus of b2 domain (res. 586) to the membrane surface should be at most 12.0 nm for optimal compatibility with the 1:1 Spike: ACE2 complex. Of the above mentioned 11 selected models in molecular docking, we found that in 6 models the distance of b2 C-terminus to the membrane surface is in between 9. 1 nm and 9. 8 nm; in one model, it is about 10.6 nm. For the other 4 models, the distance is 13.8, 13.4, 17.3 and 13.5 nm, thus eliminating these models, and leaving 7 models for consideration (Fig. 2). Overall, for all the seven models, the a2-b1-b2 Nrp1 domain containing protein is bound between the NTD (res. 13-305) and RBD (res. 318-541) of Spike protein Chain C. If the binding of the a2-b1-b2 fragment is outside of this region, it is more likely that the b2 C-terminus is too far from the membrane surface.

Based on the topological information, we would suggest that the binding of Nrp1 to Chain A and Chain B could be very stressful for Nrp1 as the b2 domain is far above the membrane in all seven models (See Fig. 5c). However, we can’t exclude the possibility that linker 2 and 3 of Nrp1 may largely straighten themselves, so that Nrp1 can still access the TNSPRRAR binding motif of Chain A and B spatially. Furthermore, in the case that Nrp1 could interact with S1 in absence of ACE2, the spatial restraint can be released. Thus, next we study the separation process when Spike protein is bound to three Nrp1. We may expect that if all three Spike protein units bind to three Nrp1, the completed separation of S2 from S1 for all three units can happen sooner with a shorter displacement of S2 domain. This would greatly facilitate the separation of S2 from S1 domain. We selected model 1 to study the separation of S2 and S1 in case of three Nrp1 binding to the three subunits of Spike protein. Fig. 6 shows the exit process of S2 from S1 with the assistance of three Nrp1s. The complete separation of S2 and S1 for Chain A, B, C occurs with a displacement of S2 domain of (6.8 nm, 9 nm, 12 nm) and (13 nm, 2.3 nm, 5.8 nm) respectively for two repeated simulations. These displacements are much shorter than the displacement of S2 domain (∼13-17 nm for all three units) in the absence of Nrp1. However, the pulling work needed to separate S2 and S1 with the support of Nrp1 is also only slightly smaller than that without the support of Nrp1: 1026±123 kcal/mol versus 1178±91 kcal/mol. Therefore, while there is not much difference in terms of the overall work needed for separation, the support of Nrp1 would alter the kinetics of S2: S1 domain. As shown in Fig. 6b, Nrp1 binding alters the energy barrier (it seems largely eliminating the second activation step) allowing earlier and faster exit of the S2 from the S1 domain.

**Figure 6:**
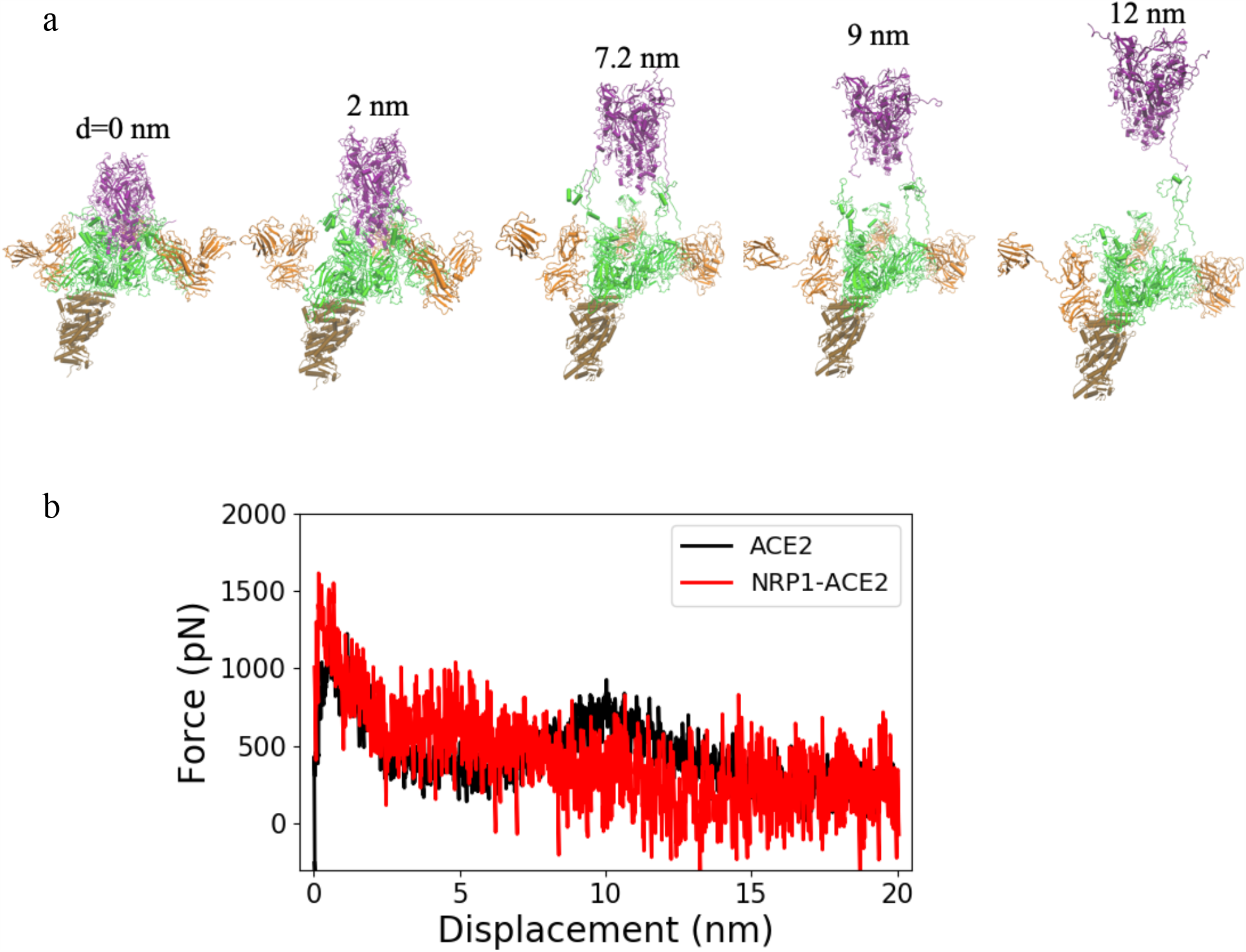
(a) Separation process of S2 and S1 domain of Chain A, B and C with three Nrp1s binding to each unit of the Spike protein trimer. The displacement of the S2 domain relative to initial position is shown. (b) Pulling force versus displacement of S2 domain with three Nrp1 binding. Averaged over 2 simulations for model 1 (red) whereas the ACE2 data is taken from Fig. 4c.

Lastly, we would emphasize that the topological or space constraints are of particularly importance in the S1 and S2 dissociation process. This basically applies to all peripheral membrane proteins interaction systems, for example Ras-Raf membrane interaction.^30,31^ The interfaces for virus Spike protein: host receptor contacts are spacially and geometrically limited. As we estimate in Fig. 5, the distance between viral and host membrane is around 30 nm. In the meantime, the exit of S2 domain out of the S1 cap-shaped domain involves a displacement of S2 domain by at least ∼5 nm (step 1). Since the membrane normal distance between viral and host membrane is limited, we suggest the separation of S2: S1 domain is unlikely merely via the membrane normal direction (Z direction), as the displacement of Spike protein at the Z direction would also triggers the displacement/deformation of viral membrane, this would result in significant energy penalty. Therefore, the tilting of Spike protein relative to membrane normal direction is very helpful, as in this case, the exit of S2 out of S1 could occur over greater distance at the lateral (X-Y) direction. In this situation, the lateral diffusion of the Spike protein relative to the ACE2 and Nrp1 receptors may provide a source of kinetic energy that drives the separation of the S2 and S1 domain. There is no evidence for the hypothesis provided here yet and future computational work may help to further unveil the mechanism.

## Summary

Neuropilin is a key binding receptor for the development and maintenance of many neural tissues and for directing axons as well as the formation of dendrites and synapses. Recently it was recognized as a novel binding receptor for SARS-CoV-2 virus. In this study, we modeled the structures of Nrp1 a2-b1-b2 binding to SRAR-CoV-2 spike protein. We analyzed the topological constraints which are generated when Nrp1 together with ACE2 binds to the Spike protein. Importantly, we studied the exit mechanism of S2 out of S1 domain with the assistance of Nrp1. We found that the Nrp1 association with Spike protein could facilitate an earlier/more probable separation of S2 and S1 domain by destabilizing the S1:S2 contacts and allowing easier dissociation of S2 N-terminal segment from this region.

## Methods

### Complex of Nrp1 a2-b1-b2 domain with Spike protein segment 678-TNSPRRAR-685

The structure of the human Nrp-1 a2-b1-b2 domain was built based on available crystal structure (PDB entry 2qqm). Missing loops in the crystal structure were added using MODELLER.^40^ The VEGF-A: Nrp-1 b1 domain complex (PDB entry 4deq) was used as a template to construct the model of the TNSPRRAR peptide bound to Nrp-1. The C-terminal residues of VEGF-A (give peptide) were mutated to give the 8 C-terminal residues of the S1 domain (above). Next, the system was simulated and relaxed with an all atom molecular dynamics simulation for 20 ns following the conventional simulation procedure (see below). The final simulation structure was extracted and used for molecular docking.

### Molecular docking of Nrp1 a2-b1-b2

#### TNSPRRAR to a Spike protein monomer

The structure of Spike protein trimer (Chain A, B, C each of res. 1-1146) was based on available crystal structure (PDB entry: 6vsb)^23^ and models (adding missing loops and residues) developed by Woo et at.^24^ The relaxed structure of the Nrp1: C-terminal octamer peptide obtained from the above simulation was treated as a ligand and was then docked to the receptor ---here the Spike protein monomer S1 domain of res. 1 to 677 (extracted from the Spike protein trimer) with ClusPro 2.0.^22^ The Ligand in this case consisted of the Nrp1 a2-b1-b2 and the Spike protein segment 678-TNSPRRAS-685. The segment (residue 678) needs to fuse with residue 677 of the receptor Spike protein. So a distance restraint of 3 Å was applied between residues 677 and 678 in the docking. 30 structures were predicted via ClusPro. Similar models, based on RMSD superposition, were merged. Next, the predicted structures were then superimposed back to the original Spike protein trimer (res. 1-1146). Structures that clash with the Spike protein trimer were excluded, reducing the number of models to 11. We also exclude models that position C-terminal of Nrp1 b2 domain (res.586) too distal from the membrane, as analyzed in the main text. This made us focus on 7 models of Nrp1 a2-b1-b2: Spike protein binding which fulfilled these criteria.

### Molecular simulation of Nrp1 a2-b1-b2

#### Spike protein trimer

We simulated the seven selected models of a complex of Nrp1 a2-b1-b2: Spike protein trimer. Standard simulation procedures were used with the NAMD/2.12 package.^41^ All proteins were solvated by TIP3P water with 150 mM NaCl. Simulation parameters were set as 2 fs for time step; thermostat at 310 K; barostat of 1 bar; and using the CHAMRM36m force field.^42^ The analysis of pair interaction and surface accessible surface area were performed with standard script within the lab.

### Exit of S2 domain from S1 domain with steered molecular dynamics

The size of S2 domain is very large, the pulling of S2 domain with explicit solvent would involve the pushing away of excessive water molecules. Since the pulling was done in limited time, these excessive water molecules would find it hard to relax themselves in a short time. This would introduce overwhelmingly high noise during the pulling. Thus, we adopted Generalized Born Implicit Solvent method.^43^ In the implicit solvent simulation, the cut-off for calculating Born radius was set as 15 Å. A cut-off of 14 Å was used for calculating the interactions between atoms. Ion concentration was set as 150 mM. For the steered molecular dynamics simulations, a time step of 1 fs was used. The S2 domain was pulled out of S1 domain with a constant velocity of 0.00001 Å per time step over 20ns of simulations. A force constant of 25 kcal/mol/Å^2^ was used for the harmonic spring that generates the force to pull S2 away from S1. During the pulling, the ACE2 protein as well as NRP1 were restrained. In order to resist the drift of S1 domain, the RBD of three Spike protein units were also restrained. The principle axis of Spike protein trimer was aligned in the Z direction. and the pulling direction was set in the Z direction. The pulling force was plotted versus the displacement of S2 domain. The total work was calculated as the cumulative work at every step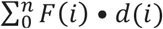 Also it should be noted that due to the applying of biased force to the system in a very limited timescale (20ns), while the SMD simulation is good way to have a look at the qualitative energy barrier for separating two molecules or studying protein unfolding, the pulling work can be significantly overestimated.^44-46^ So it should be only treated as qualitative estimation especially for S2:S1 dissociation, where S2 is a very big protein domain. For accurate free energy calculation, methods such as umbrella sampling simulations should be a better way.^47^ In addition, due to applying of biased force, the association between Nrp1 and S1 TNSPRRAR motif could be quickly broken out during the pulling. This happens for model 2, 3, 6 and especially for model 5. In order to study the role of the implication of Nrp1 binding, we exclude these simulations where Nrp1: S1 TNSPRRAR associations were not maintained during the pulling.

## Acknowledgements

We thank Prof. Tracy Tran (Rutgers’s Univ.) for insightful discussion. This work is supported by a NIH R01 grant from the National Eye Institute R01EY029169 and previous grants from NIGMS (R01GM073071 and R01GM092851) to the Buck lab.

## Author Contribution

M.B. and Z.L. conceived the project and wrote the manuscript.

## Competing Interests

The authors declare no competing interests.

## Notes

### Competing Interest Statement

The authors have declared no competing interest.

### Summary of Updates

Update the figures for better optimization (upside down S1 and S2 for the pulling snapshots).

