## supporting_information for "Neuropilin-1 Assists SARS-CoV-2 Infection by Stimulating the Separation of Spike Protein Domains S1 and S2"

**Figure S1: Residues identified as wider interface (> 30% occupancy and less than 0.6 nm) distant to the other protein.** Typically, a cut-off of 0.4 nm cut off is used to judge the interaction between two amino acids (Table in Fig. 2b). In order to include the neighboring residues at the NR1P: SPIKE interfaces, we also used a cut-off of 0.6nm, yielding interface residues shown pictorially in aligned sequence segment bars for the two proteins.

**Nrp1 domains:** a2: res. 27-141, b1: res. 147-265, b2: res. 275-424.

**Spike protein S1**: N1: res. 1-19, N2: res 20-38, N3: res. 58-90, N4: res. 210-220, N5: res. 293-318, $RBD^N$: res. 318-330, $RBD^C$: res. 525-541, C1: res. 602-610, C2: res. 621-626, C3: res. 631-644, C: res. 682-685, $S2^N$: res. 689-691

| | Nrp1 domains | Spike protein S1 |
|---|---|---|
| Model 1 | b1, b2 | N2, C2, C3, C |
| Model 2 | a2, b1 | N1, N2, N4, C |
| Model 3 | a2, b1 | N1, N2, N3, N4, $RBD^N$, $RBD^C$, C2, C, $S2^N$ |
| Model 4 | a2, b1, b2 | N1, N2, N3, N4, C3, C |
| Model 5 | b1 | N3, N4, N5, C1, C, $S2^N$ |
| Model 6 | a2, b1 | N2, N3, N4, $RBD^C$, C1,C2, C3, C, $S2^N$ |
| Model 7 | b1, b2 | N1, N2, N3, C2, C3, C |

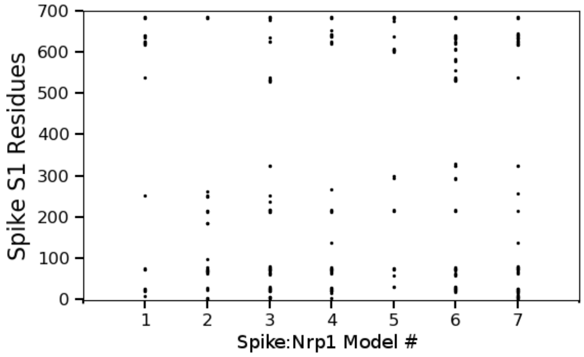


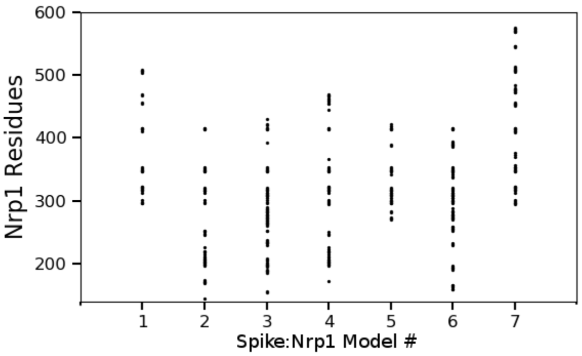
